# SpoIIDMP-driven peptidoglycan rearrangement is crucial for ribosome translocation into the spore

**DOI:** 10.1101/2024.02.12.579902

**Authors:** Olga Iwańska, Przemysław Latoch, Mariia Kovalenko, Małgorzata Lichocka, Joanna Hołówka, Remigiusz Serwa, Agata Grzybowska, Jolanta Zakrzewska-Czerwińska, Agata L. Starosta

## Abstract

In a spore-forming bacterium *Bacillus subtilis* transcription and translation are uncoupled and the translational machinery is located at the cell poles. During sporulation the cell undergoes morphological changes including asymmetric septation and chromosome translocation. However, the fate of translational machinery during sporulation has not been described. Here, using a combination of microscopic assays and mass spectrometry, we are tracking the ribosome localisation during sporulation in *B. subtilis* WT and mutants. We show that the ribosomes are associated with the asymmetric septum which is a functionally important organelle and that peptidoglycan rearrangement is essential for ribosome packing into the forespore. We also show that the feeding tube channel SpoIIIA-SpoIIQ is not required for the ribosome translocation, but is essential for maintaining the chromosome inside the spore.

**One-Sentence Summary:** Movement of ribosomes into the spore of *B. subtilis* follows chromosome transport and is precisely orchestrated in the cell.

## Introduction

The view of a bacterial cell as a disorganised collection of macromolecules enclosed by a rigid cell wall is slowly but surely going into obscurity (1). The number of proteins having discrete subcellular addresses in the bacterial cell has been increasing with examples including cell division proteins FtsZ and the Min system (2, 3) or chromosome segregation proteins FtsK and SpoIIIE in *Escherichia coli* and *Bacillus subtilis* respectively (4, 5). Interestingly, the translational machinery itself has been somewhat neglected in this type of studies, however not due to a lack of subcellular location. In vegetative cells of *B. subtilis*, the ribosomes were shown to be localised at the cell poles, away from the centrally located nucleoid and transcriptional machinery (6, 7). The functional consequences of this separation were recently described by Johnson *et al*. (8) in their seminal paper showing that transcription and translation are uncoupled in *B. subtilis*.

In response to nutrient limitation, vegetative cells of *B. subtilis* can undergo a developmental change to form dormant and resistant spores. The hallmark of sporulation is formation of an asymmetric septum which divides the cell into two compartments with differing fates: small forespore and a larger mother cell. This spectacular cell differentiation consists of precisely regulated positioning of the asymmetric septum, chromosome segregation, engulfment of the forespore by the mother cell, spore coat and cortex development and finally, mother cell lysis and spore release (9–12). In this sequence of morphological events, two processes receive considerable attention – rearrangements of the cell membrane and peptidoglycan and chromosome translocation (13–16). However, although the translational machinery including ribosomes and translation factors constitutes approximately 20% of the cell volume (17), little is known about its localisation and dynamics during sporulation, and only recently the ribosomes became objects of detailed observations in the sporulating *B. subtilis* (18).

In this work we used a combination of microscopic assays to monitor positioning of the ribosomes during sporulation in *B. subtilis*. We showed that localisation of the ribosomes correlates with the active translation sites and that the asymmetric septum plays an important role in the spatial organisation of ribosomes. We also showed that the ribosomes enter the forespore in a sequential manner, after the chromosome translocation and this is dependent on septal peptidoglycan rearrangements.

## Results

### Ribosome packing into the forespore is sequential

In this work, we have extended our initial dataset published in Iwanska *et al*. (19). Using fluorescence microscopy, we observed distribution of ribosomes during sporulation in *B. subtilis* in one-hour intervals, beginning at T0 (logarithmic growth, immediately prior to sporulation induction) to T6 (six hours post sporulation induction). We tagged the small ribosomal protein RpsB (S2) with GFP to monitor the position of ribosomes in relation to the asymmetric septum and the chromosome, stained with FM4-64 and DAPI respectively (Fig. 1A and B). At T0, mid-exponential phase, the GFP signal is localised throughout the cell, with a slight increase at the cell poles. This pattern becomes more pronounced one hour post sporulation induction, where both poles are enriched in ribosomes. However, at T2 when the asymmetric septum is being formed at approximately 20^th^ percentile of the cell’s length, we observed a drop in GFP signal in favour of DAPI at the cell pole where the forespore is developing. Approximately three hours post sporulation induction, membrane migration is mostly complete, based on the increase in FM4-64 signal at the cell pole where the spore is (Fig. 1A and B, fig. S1), and this is accompanied by the chromosome translocation resulting in forespore inflation to ∼30% of the mother cell’s length. Interestingly, we observed that during asymmetric septation and chromosome translocation (T2-T3), the ribosomes that were previously at the cell pole are restricted from the forespore rather than trapped in the divided cytoplasm. It appears that the ribosomes gather and wait at the asymmetric septum at the mother cell side and during or very shortly after the engulfment by the mother cell, the ribosomes are translocated into the spore (T4). To further investigate whether such sequential packing of the developing spore is characteristic of sporulation and applies also to transcriptional machinery, we observed the localisation of GFP-tagged β’ subunit of RNA polymerase (RpoC) during sporulation. We show that the RpoC colocalizes with the chromosome during asymmetric septation and membrane migration (T2 and T3), and is not excluded from the developing forespore prior to engulfment (T4) (Fig. 1C and D). The SpoIIIE channel strips the translocating chromosome of the DNA-binding proteins including RNA polymerase. However, as the chromosome segregation begins before asymmetric septation, a proportion of RNA polymerase is present at the cell pole prior to SpoIIIE channel formation. The increase in fluorescence intensity in the forespore results from expression of *rpoC-gfp* from the native locus, which is present in the forespore compartment at the time of polar septation (20). This is supported by the detection of *de novo* RNA synthesis in the forespore at T2 and T3, measured by incorporation and fluorescent tagging of the uridine analog, 5-ethynyluridine (EU) (21) (fig. S2). The initial expression of RpoC-GFP in the early forespore implies however the foresence of translating ribosomes in this compartment. We investigated this using an Alexa-488 labelled alkyne analog of puromycin (o-propagyl-puromycin, OPP). Puromycin terminates translation by mimicking an aminoacyl-tRNA and binding to the nascent polypeptide chain. Fluorescent labeling of OPP-bound ribosomes allows visualisation and quantification of the incorporated OPP, or in other words, of the active translation (22). As shown previously in Iwanska *et al*. (19), in the early timepoints translation localizes to the cell poles, away from the chromosome (Fig. 1E and F), which is consistent with the uncoupled transcription-translation in *B. subtilis* discussed in the seminal paper by Johnson *et al*. (8). At T2, around the time of asymmetric division, translation declines significantly which correlates with the loss of polar localisation of the ribosomes. Once the asymmetric septation is complete, translation takes place mostly at the septum on the mother cell side, and then inside the spore (T4). It should be noted however that past T4 the fluorescence signal from the spore is limited due to, most probably, spore impermeability and the results presented in Fig. 1E and F regarding late sporulation (T5 and T6) are more representative of the mother cell. Low level of translation in the forespore prior to engulfment is consistent with our observation that there is a small proportion of ribosomes present in the forming forespore and in fact, most of the ribosomes are localised at the septum on the mother cell side. Around the time of engulfment however, ribosomes are translocated into the forespore. We also followed these dynamic subcellular changes in the ribosome and RNA polymerase localisation using time-lapse microscopy (Movie S1 and S2). Based on the above observations we propose that the ribosomes are packed into the forespore sequentially, after the chromosome translocation, and that this shift is associated with membrane migration or cell wall remodelling during engulfment.

**Fig. 1.**
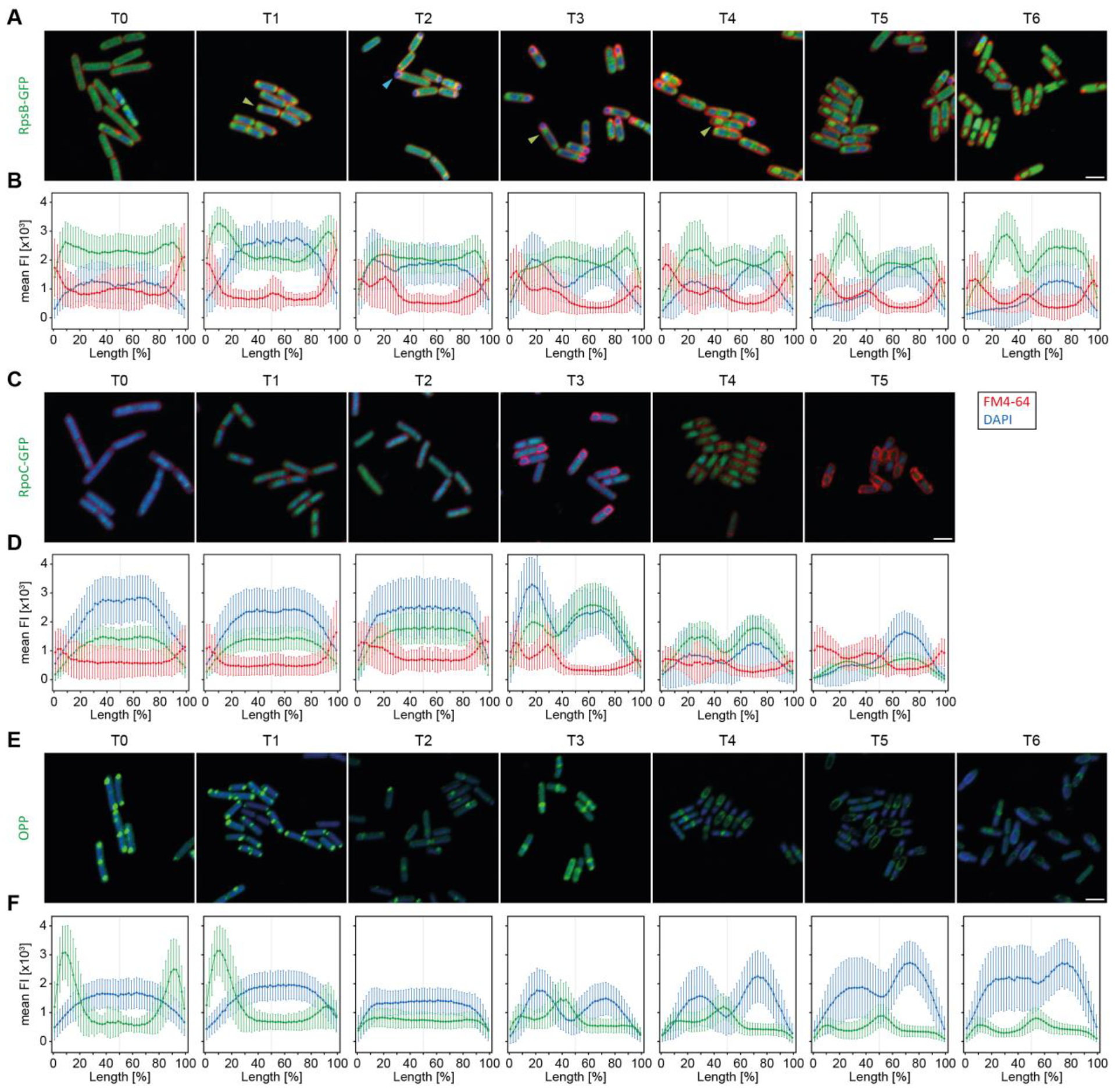
Localisation of translational machinery during sporulation in *B. subtilis* is precisely orchestrated. Sporulation process of *B. subtilis* expressing GFP-tagged ribosomes, RpsB-GFP, (**A**), or GFP-tagged RNA polymerase β’ subunit, RpoC-GFP (**C**). Cells were stained with DAPI to visualise the chromosome and FM4-64, a membrane stain, to track the asymmetric septation and sporulation progress. Samples were taken before sporulation induction (T0) and every hour of sporulation process (T1-T6). Scale bar is 2 μm. Plots of mean GFP fluorescence intensity (green) across cells show subcellular localisation of ribosomes (**B**) or RNA polymerase (**D**) during sporulation in relation to the chromosome (blue) and cell membrane and asymmetric septum (red). The cell lengths were normalised and fluorescence was measured along a line drawn across the cell long axis. (**E**) Microscopic images of *B. subtilis* WT cells treated with OPP and stained with Alexa 488 and DAPI, illustrating active translation during six hours of sporulation (T0 – T6). Blue arrowhead points to the localisation of the chromosome (DAPI) and green arrowheads point to the localisation of ribosomes (GFP). Scale bar is 2 μm. (**F**) Plots of the mean OPP-Alexa 488 fluorescence intensity (green) across the cell showing localisation and intensity of active translation during sporulation in relation to the chromosome localisation (blue). The cell lengths were normalised and fluorescence was measured along a line drawn across the cell long axis.

### Mass spectrometry indicates direct interaction of the ribosome with components of the cellular/protein machinery required for engulfment

We applied tandem mass tag-mass spectrometry (TMT-MS) analysis to determine which proteins may interact with the ribosome at the site of asymmetric septation. We collected the cells three hours post sporulation induction and performed digitonin based lysis on DSP (dithiobis(succinimidyl propionate)) cross-linked and non-cross-linked cultures, in duplicates. As controls, we used flash-frozen and pulverised cells (according to Kopik *et al*. (23)), as well as cells harvested in the logarithmic phase. Digitonin based lysis allowed for membrane solubilisation and enrichment of the lysates with membrane interacting and/or bound proteins. We then separated the 70S ribosome containing fractions using sucrose gradient centrifugation. (fig. S3). We used close to physiological salt concentration and did not prepurify the ribosomes using sucrose cushion to allow for co-migration of even transiently associated factors. Although we are aware this may result in high background, including for example large multiprotein complexes like flagellum (Fli, Flh, Flg and Hag proteins) (24), we did intend to cast a wide net in our search. The TMT-MS analysis revealed 646 proteins, including 55 ribosomal proteins (Data S1). In the cross-linked samples we observed enrichment of translation factors including IF-2 (*infB*), IF-3 (*infC*), EF-G (*fusA*), EF-Tu (*tuf*) and RRF (*frr*) indicating that we have collected actively translating ribosomes. We also observed enrichment of the cell wall and membrane associated proteins, such as components of the Min system, bacterial cytoskeletal homologs (MreB, MreC, Mbl, FtsZ, FtsA, FtsH), mother cell and forespore associated penicillin-binding proteins (PonA, DacABF), regulatory proteins (EzrA, GpsB) and specifically, components of the feeding tube channel (SpoIIIAG, SpoIIIAH and SpoIIQ) and septal peptidoglycan degradation module (SpoIIP and SpoIIB). Since both of these protein complexes are essential for engulfment, we further investigated whether either one or both may have a role in translocation of the ribosomes into the developing spore.

### Deletion of the components of SpoIIIA-SpoIIQ complex leads to forespore chromosome efflux but does not influence ribosome localization

The SpoIIIA proteins (SpoIIIAA-SpoIIIAH), together with SpoIIQ, form a multimeric channel connecting the mother cell and the forespore. The channel was proposed to function as a feeding tube and shown to maintain forespore integrity and mediate small-molecule intercellular transport (25, 26). Deletion strains of any component of the complex produce small forespores with membrane invaginations and limited transcription during late sporulation (expression of σG regulon), generally resulting in defective spore formation (27). SpoIIIAH and SpoIIQ were also identified as having additional role in membrane migration during engulfment (28). We investigated ribosome localisation during sporulation in *ΔspoIIIAH* and *ΔspoIIQ* strains in twenty minutes intervals, between two and four hours post sporulation induction, using GFP-tagged ribosomes as above. Both strains formed asymmetric septa between 2 and 2:20 hours post sporulation induction, maintaining the ribosome localisation to the mother cell side of the septum, similarly to WT cells, and failed to conclude spore engulfment by T3 as, expected (Fig. 2A and B). However, in the later stages of sporulation (T3:20 – T3:40) we detected the GFP signal at the forespore membrane vicinity, and by T4 the ribosomes were localised inside the forespore. This implies that the Q-AH complex may aid in, but is not essential for the ribosome translocation into the spore in *B. subtilis*. Interestingly, we found that the forespores containing ribosomes did not possess the chromosome in late sporulation (T4) (Fig. 2A and B, fig. S4). This, together with our data showing RpoC-GFP localisation in the spore in late sporulation (T4-T5) and rather low levels of translation in the *ΔspoIIQ* strain (fig. S5A and B) suggests chromosome translocation back into the mother cell. We therefore propose that spore deflation reported in Doan *et al*. (25) is most probably caused by an inadequate chromosome counterion transport into the spore which results in loss of turgor pressure and forespore chromosome efflux. This in turn results in limited transcription of late sporulation genes in the forespore reported by others (25, 29). A similar mechanism is seen when the chromosome fails to be transported into the spore due to defective SpoIIIE, and turgor pressure is not maintained resulting in spore deflation (18).

**Fig. 2.**
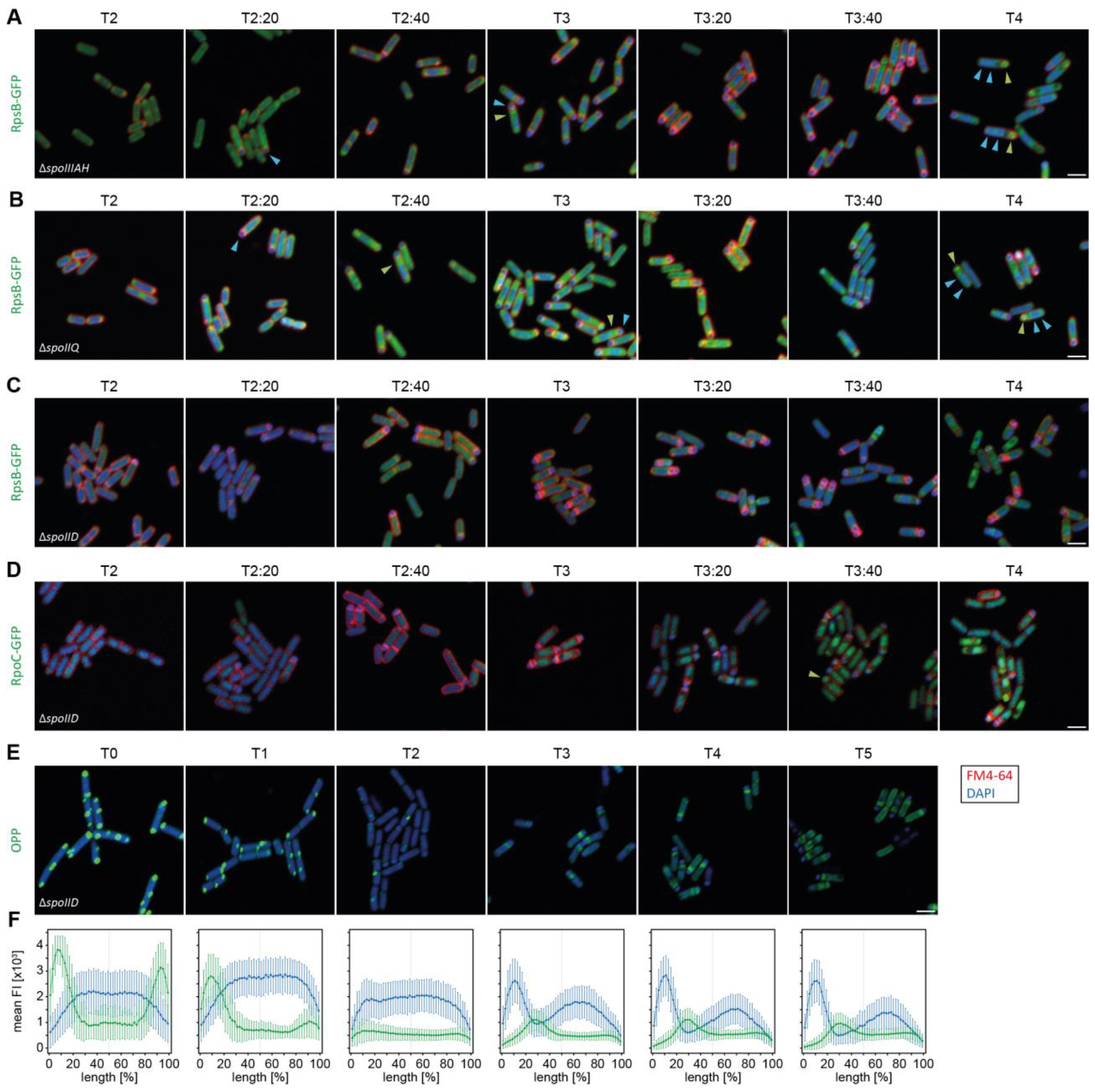
Peptidoglycan remodelling, but not membrane migration, is essential for ribosome translocation into the developing spores. (**A**), (**B**) and (**C**) *ΔspoIIIAH, ΔspoIIQ* and *ΔspoIID* strains with fluorescently tagged RpsB-GFP protein, respectively. (**D**) *ΔspoIID* strain with fluorescently labelled RNA polymerase subunit β’ (RpoC-GFP). Cells were stained with DAPI to visualise the chromosome and FM4-64, a membrane stain, to track the asymmetric septation and sporulation progress. Samples were taken every 20 minutes starting two hours post sporulation induction (T2) up to four hours post sporulation induction (T4). Scale bar is 2 μm. (**E**) Microscopic images of *ΔspoIID* strain treated with OPP and stained with Alexa 488 and DAPI, illustrating active translation during sporulation (T0 – T5). Blue arrowheads point to the localisation of a chromosome (DAPI) and green arrowheads point to the localisation of ribosomes or RNA polymerase (GFP). Scale bar is 2 μm. (**F**) Plots of the mean OPP-Alexa 488 fluorescence intensity (green) across the cell showing localisation and intensity of active translation during sporulation in relation to the chromosome localisation (blue). The cell lengths were normalised and fluorescence was measured along a line drawn across the cell long axis. Cells exhibiting the bulged membrane phenotype were selected for the analysis.

### Peptidoglycan remodelling by the SpoIIDMP is crucial for ribosome translocation into the spore

The SpoIIDMP complex partially degrades peptidoglycan at the asymmetric septum allowing for membrane migration and new peptidoglycan synthesis, and is thus essential for forespore engulfment. SpoIID deletion mutant demonstrated impaired engulfment and produced bulged septal membranes (30, 31). We therefore investigated localisation of the translational machinery (GFP-tagged ribosomes) in the *ΔspoIID* strain between two and four hours post sporulation induction. We observed asymmetric septation in the *ΔspoIID* strain between T2 and T3, followed by bulging of the septum (Fig. 2C). Similar to WT, in the *ΔspoIID* strain ribosomes are also mostly excluded from the forespore and localised at the mother cell side of the asymmetric septum (T2:40). However, unlike in WT, we did not record enrichment of the GFP signal inside the forespore at the later timepoints, suggesting that translocation of the ribosomes into the developing spore depends on peptidoglycan rearrangement. To test whether this applies also to transcriptional machinery, we observed the localisation of RpoC-GFP and recorded fluorescence signal in the forespore post asymmetric septation (Fig. 2D). We then investigated translational activity of the *ΔspoIID* strain and showed that after the asymmetric division there was negligible level of active translation in the forespores of the mutant strain (Fig. 2E and F). In the light of SpoIIIE protein stripping activity during chromosome translocation discussed above, a substantial decrease of translation together with the presence of RNA polymerase in the forespore led us to further investigate the ribosomal localisation during sporulation. We used structured illumination microscopy (SIM) to observe the localisation of ribosomes (RpsB-GFP) in the WT and *ΔspoIID* strains between one and four hours post sporulation induction in 30 minutes intervals, as well as localisation of the RNA polymerase (RpoC-GFP) in WT as a control (Fig. 3). We show that during asymmetric septation a small proportion of ribosomes is associated with the forespore membrane. In the WT strain, as the chromosome translocation concludes and as the peptidoglycan rearrangement driving spore engulfment progresses, the fluorescent signal in the spore membrane increases suggesting relocation of the ribosomes into the spore (T2 – T3). However, this is not the case for the *ΔspoIID* strain in which blockage of the peptidoglycan degradation-driven engulfment prevents ribosome translocation into the spore. As the bulged, newly synthesised membrane is devoid of the GFP signal, this implies that the cell possibly uses existing mother cell membrane proteins to anchor and transport ribosomes. This may also suggest that *de novo* ribosome biosynthesis in the developing spore is very limited if not null. Levels of RNA polymerase on the other hand are only briefly decreased in the forespore, at the time of chromosome translocation by the SpoIIIE (T2), and the fluorescence intensity and distribution quickly become similar in both mother cell and the forespore in WT.

**Fig. 3.**
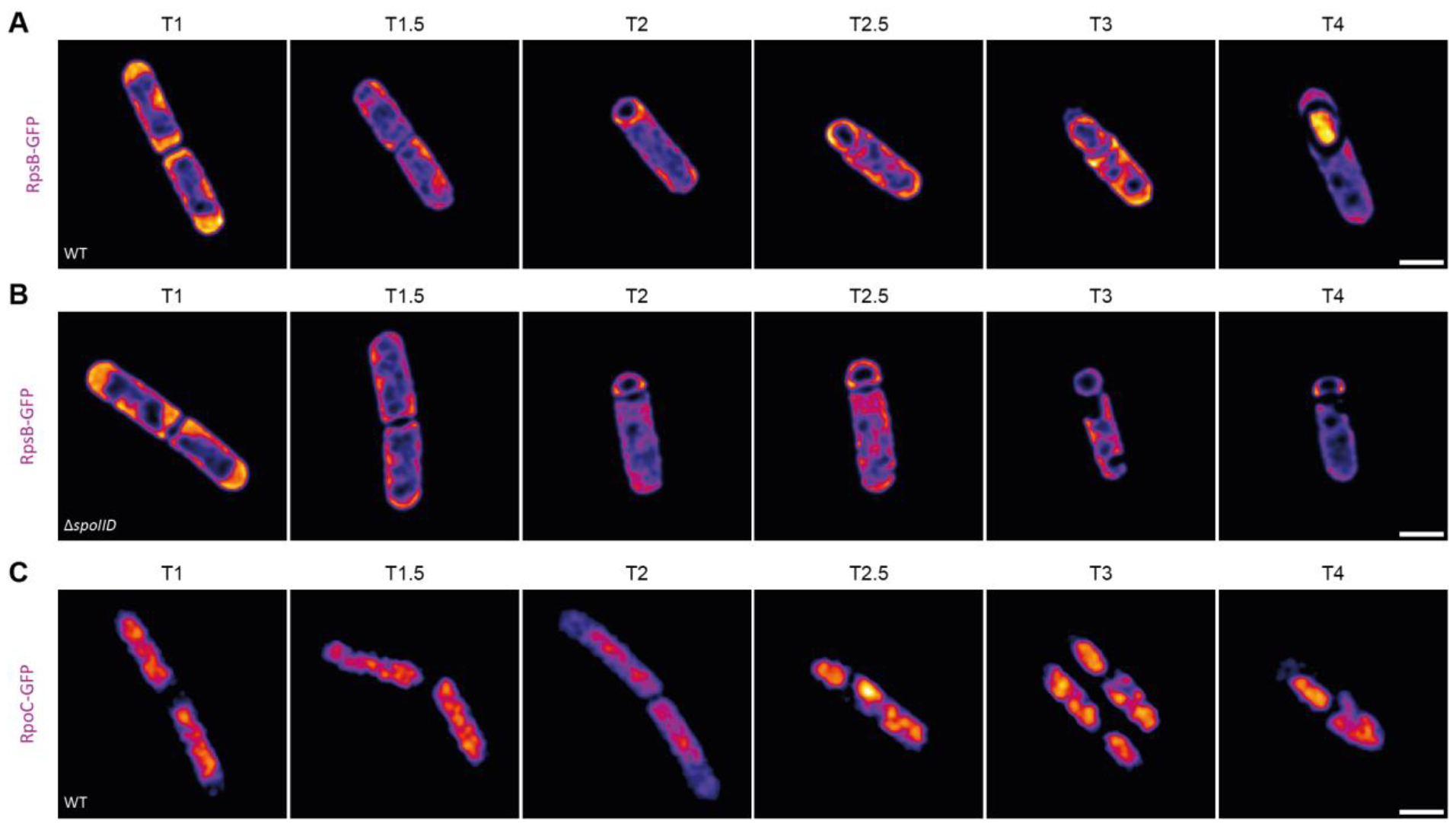
Ribosomes migrate from the mother cell to the forespore post chromosome translocation with the aid of peptidoglycan rearrangements. SIM images of WT (**A**) and *ΔspoIID* (**B**) strains with GFP-tagged ribosomes and WT strain with GFP-tagged RNA polymerase (**C**) during sporulation from T1 (one hour post sporulation induction) to T4 (four hours post sporulation induction). Images processed in ImageJ using Fire LUT represent fluorescence signal intensity (blue – low; white – high). Scale bar is 1 μm.

## Discussion

In this work, we propose a model of ribosomes translocation into the developing spore in *B. subtilis*. Contrary to the current belief, we propose that the ribosomes are not already present in the forespore and displaced to the forespore periphery during chromosome translocation (18). Rather, our data shows that ribosomes move away from the pole where the asymmetric septation will take place and only a small proportion of membrane associated ribosomes is included in the developing forespore. The majority of the ribosomes then wait at the asymmetric septum at the mother cell side where they remain translationally active. This supports the view of the asymmetric septum as a bacterial organelle serving as a hub in the cell’s developmental control, proposed by Shapiro and Losick (32). Peptidoglycan rearrangement by SpoIIDMP is crucial for ribosome packing into the spore after the chromosome translocation and we hypothesize that this transfer is mediated by mother cell membrane or cell wall proteins. Once the chromosome becomes dehydrated, ribosomes are released from the spore membrane and move towards the centrally located chromosome. The proposed model implies two things – (I) ribosomes can be transported to different subcellular locations during development and the cell most likely employs homologs of cytoskeletal proteins for this; (II) the cues for localisation of asymmetric septation may be translational and also based on the sites of ribosome location. Whether membrane targeting and membrane association of the ribosomes is via signal recognition particles, short nascent peptides (33) or by direct binding with the membrane or cell wall components (perhaps lipid II mediated?), remains to be answered.

We postulate that the SpoIIDMP role in ribosomes translocation into the forespore can be underlined by the evolutionary conservation of these proteins in the sporulation gene set of *Firmicutes* (34) and can suggest that ribosome transport may be widespread and universal in spore forming bacteria. Since we did not observe ribosome biogenesis factors in the MS data nor the GFP signal in the bulged membrane of *ΔspoIID*, this suggests that ribosomes are not synthesised *de novo* in the forespore and ribosomal transport from the mother cell to the developing spore is in fact required for successful sporulation. SpoIIIA-SpoIIQ feeding tube channel is another example of a conserved complex essential for engulfment in sporulating bacteria. Although our data shows that it is not necessary for ribosome transport into the forespore, we do not exclude the possibility that it may play an indirect role. However, we propose that the function of the feeding tube channel, apart from its role in engulfment, is transport of small molecules contributing to establishing turgor and maintaining the chromosome inside the spore. It can be speculated that perhaps such function in chromosome upkeep, together with the complex structure and transmembrane fusing (35), may be reminiscent of a primal nuclear pore complex-like structure and an evolutionary step leading to formation of the nucleus.

Collectively, our results contribute to the changing idea of the bacterial cell as a spatially organised system and place the translational machinery in the context of dynamic subcellular localisation during sporulation.

## Author contributions

Conceptualization: ALS, JZC

Investigation: OI, MK, ML, JH, RS, AG

Visualization: PL

Funding acquisition: ALS

Writing – original draft: OI

Writing – review & editing: ALS, PL

## Data and materials availability

The mass spectrometry proteomics data have been deposited to the ProteomeXchange Consortium (36) via the PRIDE (37) partner repository with the dataset identifier PXD047497.

## Materials and Methods

### Strains and growth conditions

*B. subtilis* 168 strain was used as a wild type and the remaining strains are derivatives thereof, as listed in Table S1A. The strains were grown overnight in LB medium at 30 °C with shaking then diluted to OD_600_=0.1 in CH medium (38) and grown until OD_600_ reached 0.5 – 0.6, at 37 °C with shaking. Sporulation was induced by medium exchange to sporulation medium as described by Sterlini and Mandelstam (39), except the cells were harvested by filtration and filters were transferred into the culture flasks. The sporulation medium was supplemented with 3% v/v of the culture in CH medium at OD_600_=0.5-0.6 to promote sporulation.

### Strains construction

The primers and plasmids used in this study are listed in Supplementary Tables S1B and C, respectively.

*B. subtilis* single deletion strains were purchased from Bacillus Genetic Stock Center (BGSC, www.bgsc.org) (40).

For strains with the GFP tag, the RpsB ribosomal protein or β’ subunit of RNA polymerase, RpoC, were tagged with GFP at the C termini. The fusion was performed by a double cross-over and integrated into the chromosome in the native locus. The tag was introduced by transforming the competent *B. subtilis* cells (41) with a linear DNA construct prepared by overlap PCR method (42). The approximately 5,5 kb DNA construct consisted of two genomic regions 2kb upstream and downstream of the STOP codon of *rpsB or rpoC* genes amplified from the genomic DNA of *B. subtilis* using dedicated UPFOR/UPREV and DOWNFOR/DOWNREV primers, and GFP tag and spectinomycin resistance cassette were amplified from pSHP2 plasmid with MID1FOR/MID2REV primers introducing appropriate flanking regions (Tables S1B and C). The STOP codons of *rpsB* and *rpoC* were omitted. The transformants were selected on nutrient agar plates with spectinomycin and the results of transformations were verified by PCR and visually (expression of GFP).

### Confocal microscopy

Confocal microscopy was carried out with an inverted confocal system Nikon C1. Images were taken with Plan-Apo VC 100x/1.40 oil immersion objective. GFP/Alexa 488 were excited at 488 nm, DAPI at 408 nm, FM4-64 at 543 nm. The fluorescence signals were collected using filter sets: 515/30 nm for GFP/Alexa 488, 480/40 nm for DAPI and 610LP nm for FM4-64. Imaging of specimens having more than one fluorophore was performed in a sequential scan mode to prevent bleed-through of signal. *B. subtilis* strains were grown and sporulation was induced as described above. Aliquots of the cultures were sampled every hour for six hours into sporulation, beginning at time T0 – prior to sporulation induction. Cells were immobilised on 1% agarose pads prepared with 1.0 × 1.0-cm GeneFrames (Thermo Fisher Scientific). Unless stated otherwise, the cells were visualised with SynaptoRed dye at a final concentration of 10 μg ml^-1^ and 4′,6-diamidino-2-phenylindole (DAPI) dye at a final concentration of 5 μg ml^-1^. For translation visualisation, cells were incubated with O-propargyl-puromycin (OPP) using Click-iT® Plus OPP Alexa Fluor® 488 Protein Synthesis Assay Kit (Invitrogen) according to the manufacturer’s guidelines. Briefly, cells we incubated with 13 μM OPP for 30 min at 37 °C with shaking. Cells were then fixed with 3.7% formaldehyde and permeabilized with 0.1% Triton X-100. Fluorescent labelling was performed for 30 min with Alexa Fluor 488 reaction cocktail and then, cells were stained with DAPI.

### Time Lapse Microfluidics Microscopy (TLMM)

TLMM experiments were performed using a Delta Vision Elite inverted microscope equipped with a 100×/1.4 Oil objective. *B. subtilis* strains were grown and sporulation was induced as described above. Immediately after sporulation induction cells were loaded into the ONIX observation chamber (B04A-03-5PK CellASIC ONIX plates for bacteria cells) and further cultured in sporulation medium with addition of 0.57 – 3% of the rich medium. Images were recorded at 5 min intervals for 6 h (35 ms exposure time, 10% transmittance on transmitted light channel, 15 ms exposure time and 10% of transmittance on EGFP channel). The temperature during the experiment was maintained at 37°C.

### Lattice SIM Imaging

Strains were grown and sporulation was induced as described above. Cultures were sampled hourly beginning with T1 (one hour post sporulation induction), to T4 (four hours post sporulation induction). Lattice SIM Imaging was performed using an Elyra 7 (Zeiss) inverted microscope equipped with an sCMOS 4.2 CL HS camera and an alpha Plan-Apochromat 100×/1.46 Oil DIC M27 objective with an Optovar 1.6x magnification changer. Fluorescence was excited with 488 nm laser (100 mW), the signals were observed through multiple beam splitter (405/488/561/641 nm) and laser block filters (405/488/561/641 nm). Cells were immobilised on 1% agarose pads (1.0 × 1.0-cm GeneFrames; Thermo Fisher Scientific). Cells were illuminated with 488 nm laser with 0.9% intensity and 150 ms exposure time and with 0.5% intensity and 100 ms exposure time for respectively RpsB-GFP and RpoC-GFP fusions. Each Z-plane was illuminated in Lattice-SIM mode comprising 13 phases. Image reconstruction was performed with the ZEN 3.0 SR software (Zeiss) using SIM^2^ reconstruction algorithm with standard parameters and 3D models of the obtained data were prepared utilizing the VolumeViewer plugin (Fiji).

### Microscopy data analysis

Images were analysed using ImageJ (43) and graphs were prepared in R (44). For fluorescence localization measurements, a 1p line was drawn across the cells (region of interest, ROI, n>84 for GFP fluorescence, n>186 for OPP-Alexa 488 fluorescence). The cell lengths were normalised to 100% and the fluorescence intensities across the line were recorded and visualised. To enhance the visualization of the green channel intensity (OPP and EU), the minimum and maximum values of the green channel histogram were scaled to a range of 0-2500 using ImageJ “Brightness/Contrast…” option.

### Ribosome purification and mass spectrometry

Ribosomes were purified from *B. subtilis* WT cultures harvested 3h post sporulation induction, in duplicates. Before harvesting, cultures were treated with 0.3 mM chloramphenicol. Cross-linking was performed using 2.5 mM DSP for 30s, followed by quenching with 50 mM Tris for 30s. The cross-linked cultures were collected by centrifugation, 10 min, 4000 rpm, 4°C. Otherwise, cells were collected by filtration and flash frozen in liquid nitrogen. In case of detergent based lysis, frozen cells were incubated in lysis buffer (20 mM Tris pH 7.6, 10 mM MgAcet, 150 mM KAcet and 1 mM chloramphenicol), with 1% digitonin, 1mg/ml lysozyme and 100 U/ml Dnase I for 30 min at 30°C. Lysis by grinding was performed with mortar and pestle with addition of aluminium oxide, in lysis buffer without digitonin (20 mM TRIS pH 7.6, 10 mM MgAcet, 150 mM KAcet, 0.4% TRITON X-100, 1 mM chloramphenicol, 100U/ml Dnase I). As a control, cultures in logarithmic growth phase were also harvested. These were grown in CH medium at 37 °C with shaking and harvested when OD_600_ = 0.8. Lysed cells were then centrifuged and debris-free lysates were collected. Concentration of RNA was measured spectrophotometrically. Linear sucrose gradients (5% - 30% w/v, 12.5ml) were prepared in polysome buffer (20 mM Tris, 10 mM MgAcet, 5 mM KAcet,100 mM NH_4_Cl) with 0.1% addition of digitonin for the samples lysed by detergent based lysis, in polypropylene ultracentrifuge tubes. The same amounts of samples (700 μg of total RNA) were layered on top of the sucrose gradients and centrifuged in a Thermo Scientific Sorval WX Ultra Series ultracentrifuge for three hours at 28 000 rpm, 4°C using a Thermo Scientific Th-641 swinging bucket rotor. Polysome profiles were recorded at 254 nm and gradients were fractionated into 500 μl fractions using BioComp’s Piston Gradient Fractionator™. Polysome profiles were analysed and fractions corresponding to the monosome were sent for mass spectrometry analysis.

The collected samples were subjected to chloroform/methanol precipitation and the resulting protein pellets were washed with methanol. The pellets were air-dried for 10 min and resuspended in 100 mM HEPES pH 8.0. TCEP (final conc. 10 mM) and chloroacetamide (final conc. 40 mM). Trypsin was added in a 1:100 enzyme-to-protein ratio and incubated overnight at 37°. The digestion was terminated by the addition of trifluoroacetic acid (TFA, final conc. 1%). The resulting peptides were labelled using an on-(StageTip) column TMT labelling protocol (45). Peptides were eluted with 60 μl 60% acetonitrile/0.1% formic acid in water. Equal volumes of each sample were pooled into two TMT12plex samples and dried using a SpeedVac concentrator. Peptides were then loaded on StageTip columns and washed with 0.1% TEA/5%ACN, eluted with 0.1% TEA/50%ACN, and dried using a SpeedVac concentrator. Prior to LC-MS measurement, the peptide fractions were dissolved in 0.1% TFA, 2% acetonitrile in water. Chromatographic separation was performed on an Easy-Spray Acclaim PepMap column 50 cm long × 75 μm inner diameter (Thermo Fisher Scientific) at 55°C by applying 180 min acetonitrile gradients in 0.1% aqueous formic acid at a flow rate of 300 nl/min. An UltiMate 3000 nano-LC system was coupled to a Q Exactive HF-X mass spectrometer via an easy-spray source (all Thermo Fisher Scientific). The Q Exactive HF-X was operated in TMT mode with survey scans acquired at a resolution of 60,000 at m/z 200. Up to 15 of the most abundant isotope patterns with charges 2-5 from the survey scan were selected with an isolation window of 0.7 m/z and fragmented by higher-energy collision dissociation (HCD) with normalized collision energies of 32, while the dynamic exclusion was set to 35 s. The maximum ion injection times for the survey scan and the MS/MS scans (acquired with a resolution of 45,000 at m/z 200) were 50 and 96 ms, respectively. The ion target value for MS was set to 3e6 and for MS/MS to 1e5, and the minimum AGC target was set to 1e3. Data was processed with MaxQuant v. 1.6.17.0 (46), and peptides were identified from the MS/MS spectra searched against a modified Uniprot Bacillus Subtilis reference proteome (UP000001570 was extended by four entries: protein RpsN1 sequence MAKKSMIAKQQRTPKFKVQEYTRCERCGRPHSVIRKFKLCRICFRELAYKGQIPGVKKA SW; protein RpsN2 sequence MAKKSKVAKELKRQQLVEQYAGIRRELKEKGDYEALSKLPRDSAPGRLHN RCMVTGRPRAYMRKFKMSRIAFRELAHKGQIPGVKKASW; protein RpmG2 sequence MRKKITLACKTCGNRNYTTMKSSASAAERLEVKKYCSTCNSHTAHLETK; protein RpmG3 sequence MRVNVTLACTETGDRNYITTKNKRTNPDRLELKKYSPRLKKYTLHRETK) using the built-in Andromeda search engine. Reporter ion MS2-based quantification was applied with reporter mass tolerance = 0.003 Da and min. reporter PIF = 0.75. Normalization was selected, weighted ratio to all TMT channels. Cysteine carbamidomethylation was set as a fixed modification and methionine oxidation as well as glutamine/asparagine deamination were set as variable modifications. For in silico digests of the reference proteome, cleavages of arginine or lysine followed by any amino acid were allowed (trypsin/P), and up to two missed cleavages were allowed. The false discovery rate (FDR) was set to 0.01 for peptides, proteins, and sites. A match between runs was enabled. Other parameters were used as pre-set in the software. Unique and razor peptides were used for quantification enabling protein grouping (razor peptides are the peptides uniquely assigned to protein groups and not to individual proteins). Reporter intensity corrected values (values) for protein groups were loaded into Perseus v. 1.6.10.0 (47). Standard filtering steps were applied to clean up the dataset: reverse (matched to decoy database), only identified by site, and potential contaminant (from a list of commonly occurring contaminants included in MaxQuant) protein groups were removed. The values were log2 transformed and proteins groups with at least one valid value across all 11 sample groups were kept. The values were then normalized to the ribosomal protein content of each sample (within each sample a median value of 54 detected ribosomal proteins was subtracted from the values of non-ribosomal proteins). Log2 fold changes for 10 experimental groups vs Ctrl were calculated. The mass spectrometry proteomics data have been deposited to the ProteomeXchange Consortium (36) via the PRIDE (37) partner repository with the dataset identifier PXD047497.

## References

1. D. Z. Rudner, R. Losick, Protein Subcellular Localization in Bacteria. Cold Spring Harb. Perspect. Biol. 2, a000307 (2010).

2. K. D. Whitley, et al., FtsZ treadmilling is essential for Z-ring condensation and septal constriction initiation in Bacillus subtilis cell division. Nat. Commun. 2021 121 12, 1–13 (2021).

3. H. Feddersen, L. Würthner, E. Frey, M. Bramkamp, Dynamics of the bacillus subtilis min system. MBio 12 (2021).

4. H. Chan, A. M. T. Mohamed, I. Grainge, C. D. A. Rodrigues, FtsK and SpoIIIE, coordinators of chromosome segregation and envelope remodeling in bacteria. Trends Microbiol. 30, 480–494 (2022).

5. J. Bath, Ling Juan Wu, J. Errington, J. C. Wang, Role of Bacillus subtilis SpoIIIE in DNA transport across the mother cell-prespore division septum. Science (80-.). 290, 995–997 (2000).

6. P. J. Lewis, S. D. Thaker, J. Errington, Compartmentalization of transcription and translation in Bacillus subtilis. EMBO J. 19, 710 (2000).

7. J. Mascarenhas, M. H. W. Weber, P. L. Graumann, Specific polar localization of ribosomes in Bacillus subtilis depends on active transcription. EMBO Rep. 2, 685 (2001).

8. G. E. Johnson, J. B. Lalanne, M. L. Peters, G. W. Li, Functionally uncoupled transcription–translation in Bacillus subtilis. Nat. 2020 5857823 585, 124–128 (2020).

9. J. Errington, Regulation of endospore formation in Bacillus subtilis. Nat. Rev. Microbiol. 2003 12 1, 117–126 (2003).

10. P. J. Piggot, D. W. Hilbert, Sporulation of Bacillus subtilis. Curr. Opin. Microbiol. 7, 579–586 (2004).

11. D. Higgins, J. Dworkin, Recent progress in Bacillus subtilis sporulation. FEMS Microbiol. Rev. 36, 131–148 (2012).

12. K. Khanna, J. Lopez-Garrido, K. Pogliano, Shaping an Endospore: Architectural Transformations During Bacillus subtilis Sporulation. 10.1146/annurev-micro-022520-074650 74, 361–386 (2020).

13. T. C. Fleming, et al., Dynamic SpoIIIE assembly mediates septal membrane fission during Bacillus subtilis sporulation. Genes Dev. 24, 1160–1172 (2010).

14. K. Khanna, et al., The molecular architecture of engulfment during Bacillus subtilis sporulation. Elife 8 (2019).

15. A. Mohamed, et al., Chromosome Segregation and Peptidoglycan Remodeling Are Coordinated at a Highly Stabilized Septal Pore to Maintain Bacterial Spore Development. Dev. Cell 56, 36–51.e5 (2021).

16. K. Cramer, S. C. M. Reinhardt, A. Auer, J. Y. Shin, R. Jungmann, Comparing divisome organization between vegetative and sporulating Bacillus subtilis at the nanoscale using DNA-PAINT. Sci. Adv. 10, eadk5847 (2024).

17. E. Bosdriesz, D. Molenaar, B. Teusink, F. J. Bruggeman, How fast-growing bacteria robustly tune their ribosome concentration to approximate growth-rate maximization. FEBS J. 282, 2029–2044 (2015).

18. J. Lopez-Garrido, et al., Chromosome Translocation Inflates Bacillus Forespores and Impacts Cellular Morphology. Cell 172, 758–770.e14 (2018).

19. O. Iwańska, et al., Translation in Bacillus subtilis is spatially and temporally coordinated during sporulation. bioRxiv, 2023.12.06.569898 (2023).

20. K. A. Marquis, et al., SpoIIIE strips proteins off the DNA during chromosome translocation. Genes Dev. 22, 1786–1795 (2008).

21. C. Y. Jao, A. Salic, Exploring RNA transcription and turnover in vivo by using click chemistry. Proc. Natl. Acad. Sci. U. S. A. 105, 15779–15784 (2008).

22. J. Liu, Y. Xu, D. Stoleru, A. Salic, Imaging protein synthesis in cells and tissues with an alkyne analog of puromycin. Proc. Natl. Acad. Sci. U. S. A. 109, 413–418 (2012).

23. N. Kopik, O. Chrobak, P. Latoch, M. Kovalenko, A. L. Starosta, RIBO-seq in Bacteria: a Sample Collection and Library Preparation Protocol for NGS Sequencing. JoVE (Journal Vis. Exp. 2021, e62544 (2021).

24. S. Maki-Yonekura, K. Yonekura, K. Namba, Conformational change of flagellin for polymorphic supercoiling of the flagellar filament. Nat. Struct. Mol. Biol. 2010 174 17, 417–422 (2010).

25. T. Doan, et al., Novel Secretion Apparatus Maintains Spore Integrity and Developmental Gene Expression in Bacillus subtilis. PLOS Genet. 5, e1000566 (2009).

26. E. P. Riley, C. Schwarz, A. I. Derman, J. Lopez-Garrido, Milestones in Bacillus subtilis sporulation research. Microb. Cell 8, 1 (2020).

27. C. Morlot, C. D. A. Rodrigues, The New Kid on the Block: A Specialized Secretion System during Bacterial Sporulation. Trends Microbiol. 26, 663–676 (2018).

28. A. Rubio, K. Pogliano, Septal localization of forespore membrane proteins during engulfment in Bacillus subtilis. EMBO J. 23, 1636–1646 (2004).

29. A. H. Camp, R. Losick, A feeding tube model for activation of a cell-specific transcription factor during sporulation in Bacillus subtilis. Genes Dev. 23, 1014–1024 (2009).

30. C. Morlot, T. Uehara, K. A. Marquis, T. G. Bernhardt, D. Z. Rudner, A highly coordinated cell wall degradation machine governs spore morphogenesis in Bacillus subtilis. Genes Dev. 24, 411–422 (2010).

31. J. Gutierrez, R. Smith, K. Pogliano, SpoIID-mediated peptidoglycan degradation is required throughout engulfment during Bacillus subtilis sporulation. J. Bacteriol. 192, 3174–3186 (2010).

32. L. Shapiro, R. Losick, Dynamic Spatial Regulation in the Bacterial Cell. Cell 100, 89–98 (2000).

33. T. Bornemann, J. Jöckel, M. V. Rodnina, W. Wintermeyer, Signal sequence–independent membrane targeting of ribosomes containing short nascent peptides within the exit tunnel. Nat. Struct. Mol. Biol. 2008 155 15, 494–499 (2008).

34. M. Y. Galperin, N. Yutin, Y. I. Wolf, R. V. Alvarez, E. V. Koonin, Conservation and Evolution of the Sporulation Gene Set in Diverse Members of the Firmicutes. J. Bacteriol. 204 (2022).

35. M. Beck, E. Hurt, The nuclear pore complex: understanding its function through structural insight. Nat. Rev. Mol. Cell Biol. 2016 182 18, 73–89 (2016).

36. E. W. Deutsch, et al., The ProteomeXchange consortium at 10 years: 2023 update. Nucleic Acids Res. 51, D1539–D1548 (2023).

37. Y. Perez-Riverol, et al., The PRIDE database resources in 2022: a hub for mass spectrometry-based proteomics evidences. Nucleic Acids Res. 50, D543–D552 (2022).

38. M. E. Sharpe, P. M. Hauser, R. G. Sharpe, J. Errington, Bacillus subtilis cell cycle as studied by fluorescence microscopy: Constancy of cell length at initiation of DNA replication and evidence for active nucleoid partitioning. J. Bacteriol. 180, 547–555 (1998).

39. J. M. Sterlini, J. Mandelstam, Commitment to sporulation in Bacillus subtilis and its relationship to development of actinomycin resistance. Biochem. J. 113, 29–37 (1969).

40. B. M. Koo, et al., Construction and Analysis of Two Genome-Scale Deletion Libraries for Bacillus subtilis. Cell Syst. 4, 291–305.e7 (2017).

41. H. F. Jenkinson, Altered arrangement of proteins in the spore coat of a germination mutant of Bacillus subtilis. J. Gen. Microbiol. 129, 1945–1958 (1983).

42. M. Forloni, A. Y. Liu, N. Wajapeyee, Creating Insertions or Deletions Using Overlap Extension Polymerase Chain Reaction (PCR) Mutagenesis. Cold Spring Harb. Protoc. 2018, pdb.prot097758 (2018).

43. J. Schindelin, et al., Fiji: an open-source platform for biological-image analysis. Nat. Methods 2012 97 9, 676–682 (2012).

44. R Core Team, R: A language and environment for statistical computing. https://www.R-project.org.

45. S. A. Myers, et al., Streamlined Protocol for Deep Proteomic Profiling of FAC-sorted Cells and Its Application to Freshly Isolated Murine Immune Cells*. Mol. Cell. Proteomics 18, 995a–1009 (2019).

46. S. Tyanova, T. Temu, J. Cox, The MaxQuant computational platform for mass spectrometry-based shotgun proteomics. Nat. Protoc. 2016 1112 11, 2301–2319 (2016).

47. S. Tyanova, et al., The Perseus computational platform for comprehensive analysis of (prote)omics data. Nat. Methods 2016 139 13, 731–740 (2016).

48. V. Murina, et al., ABCF ATPases Involved in Protein Synthesis, Ribosome Assembly and Antibiotic Resistance: Structural and Functional Diversification across the Tree of Life. J. Mol. Biol. 431, 3568–3590 (2019).

