## Supplementary material for "SpoIIDMP-driven peptidoglycan rearrangement is crucial for ribosome translocation into the spore": Fig. S1, Fig. S2, Fig. S3, Fig. S4, Fig. S5, Table S1

**The PDF file includes:**

Figs. S1 to S5
Table S1
References and Notes

**Other Supplementary Materials for this manuscript include the following:**

Movies S1 and S2
Data S1

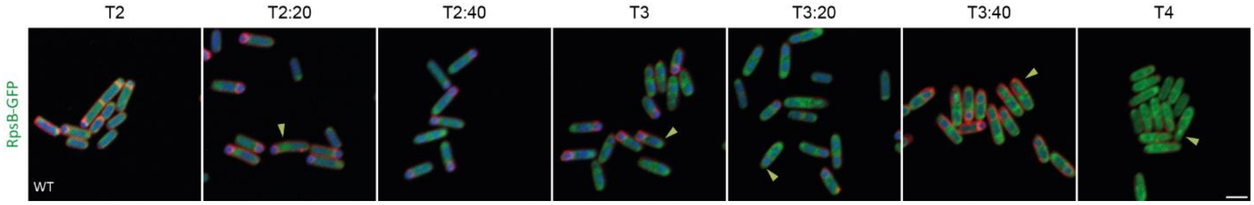


**Fig. S1. Sporulation process of *B. subtilis* expressing GFP-tagged ribosomes RpsB-GFP (green).**

Cells were stained with DAPI (blue) to visualise the chromosome and FM4-64 (red), a membrane stain, to track the asymmetric septation and sporulation progress. Samples were taken two hours after sporulation induction (T2) and every twenty minutes of the sporulation process (T2:20-T4). Scale bar is 2 μm.

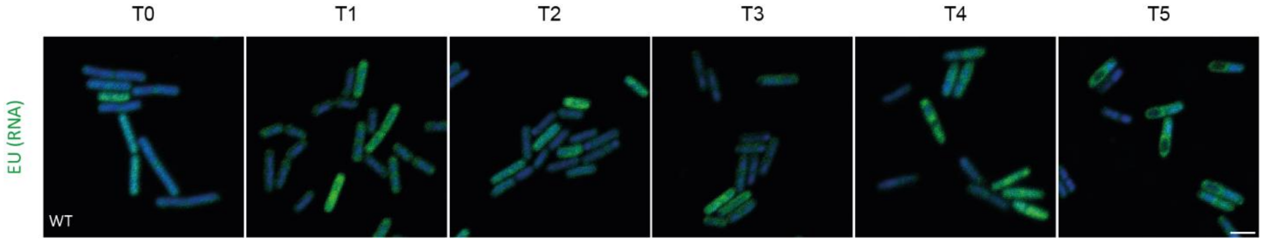


**Fig. S2. Localisation of *de novo* RNA synthesis during sporulation in *B. subtilis*.**

Cells were stained with DAPI to visualise the chromosome (blue), and *de novo* RNA synthesis was observed by incorporation and fluorescent tagging of the uridine analog, 5-ethynyluridine (EU, green). Samples were taken before sporulation induction (T0) and every hour of the sporulation process (T1-T5). The scale bar is 2 μm.

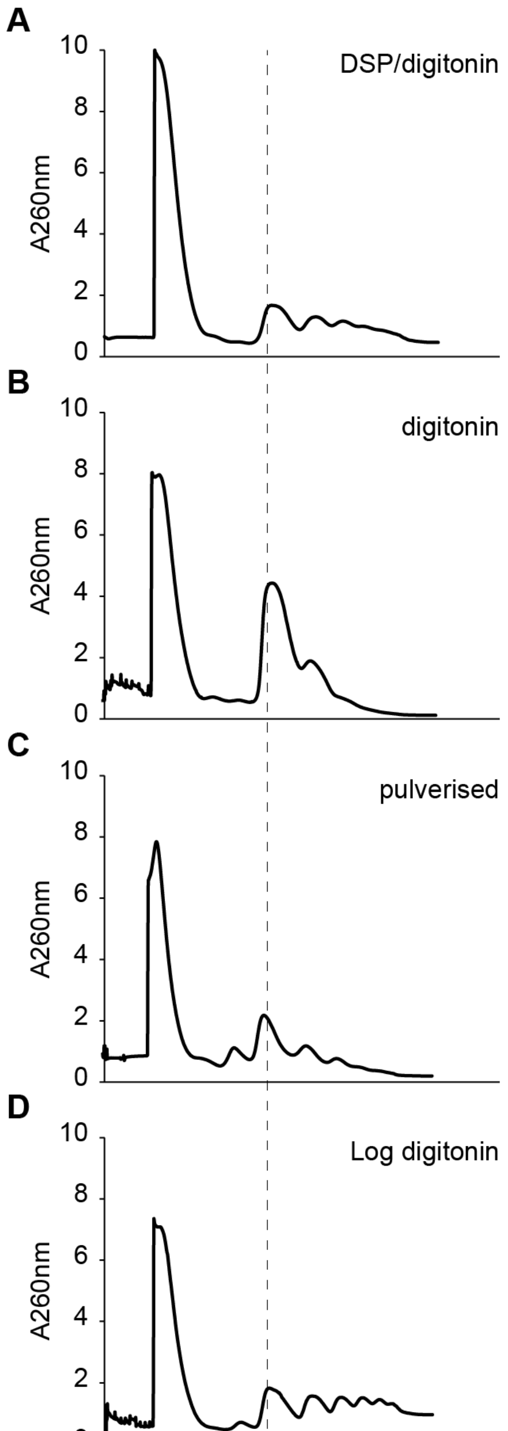


**Fig. S3. Polysome profiles of sporulating WT cells.**

Lysates were prepared by (**A**) DSP cross-linking and digitonin-based lysis, (**B**) digitonin-based lysis, (**C**) flash-frozen and pulverised with mortar and pestle, (**D**) culture harvested in the logarithmic phase and lysed with digitonin.

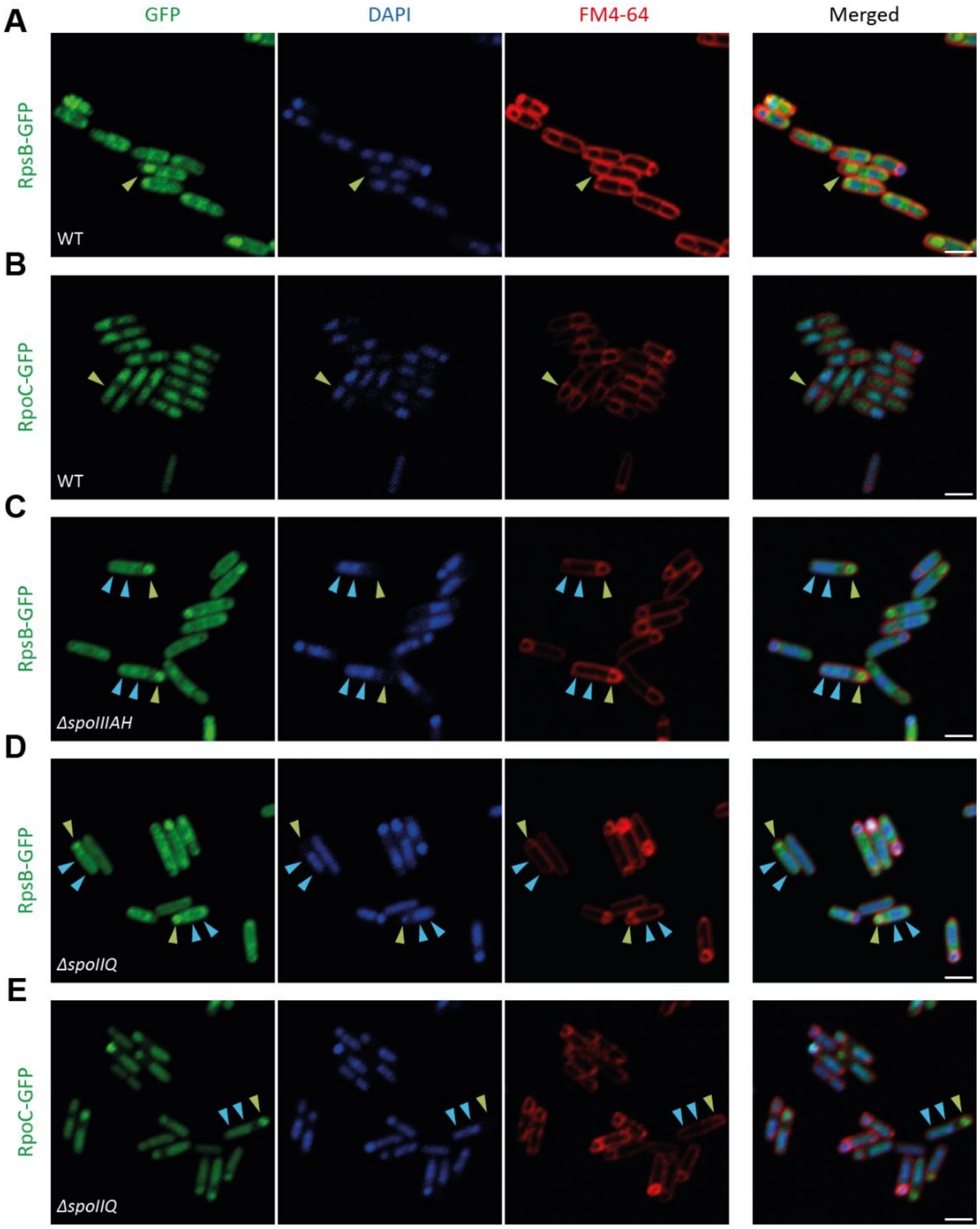


**Fig. S4. Chromosome is translocated back into the mother cell in the mutants lacking components of the feeding tube channel - ΔspoIIQ and ΔspoIIIAH.**

Single channel view for (**A**) WT, (**C**) *ΔspoIIIAH* and (**D**) *ΔspoIIQ* strains with fluorescently labelled ribosomes (RpsB-GFP) collected at T4 (four hours post sporulation induction). (**B**) WT and (**E**) *ΔspoIIQ* strains with fluorescently labelled RNA polymerase (RpoC-GFP), collected at T4. Green arrowheads point to the localisation of the ribosomes, blue arrowheads point to the localisation of the chromosome. Double blue arrowheads show the localisation of forespore chromosome that was translocated into the mother cell. Scale bar is 2 µm.

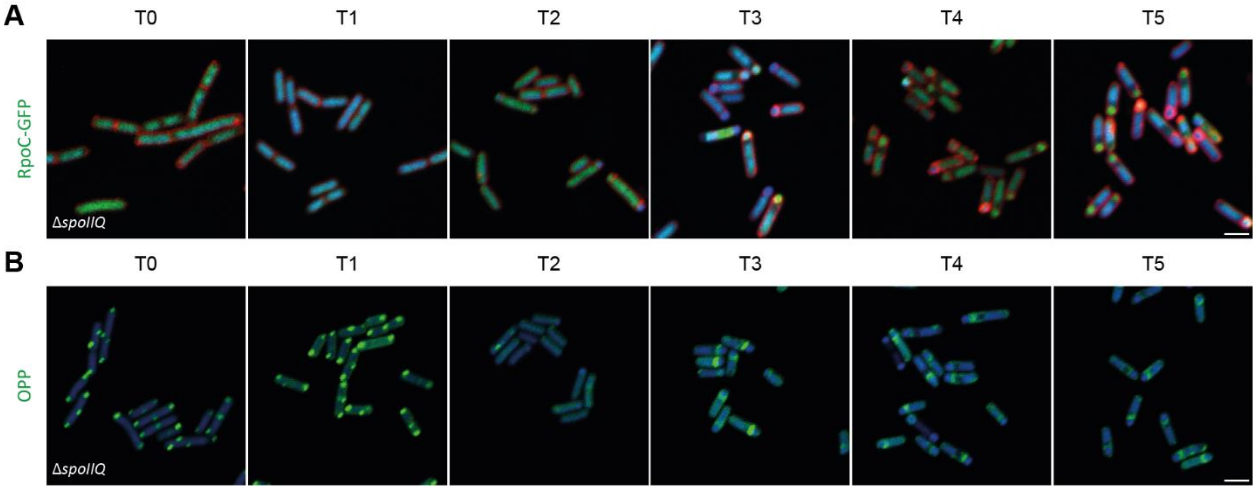


**Fig. S5. Spatial coordination of translational in *ΔspoIIQ* strain during sporulation**

(**A**) Microscopic images depict the localization of GFP-tagged β' subunit of RNA polymerase (RpoC, green) in *ΔspoIIQ* strain during different stages of sporulation. The cells were stained with DAPI to highlight the chromosome (blue). (**B**) Microscopic images of *B. subtilis ΔspoIIQ* cells treated with OPP and stained with Alexa 488 (green) and DAPI (blue). Samples were collected before sporulation induction (T0) and at hourly intervals throughout the first five hours of the sporulation process (T1-T5). The scale bar represents 2 μm.

**Table S1. List of (A) strains, (B) primers and (C) plasmids used in this study.**

**A. Strains**

| Strain | Description | Reference |
|---|---|---|
| WT | *Bacillus subtilis subsp. subtilis trpC2* | Koo *et al.* (40) |
| WT RpsB-GFP | *Bacillus subtilis subsp. subtilis trpC2 rpsB:gfp SpR* | Iwanska *et al*. 2023 (19) |
| WT RpoC-GFP | *Bacillus subtilis subsp. subtilis trpC2 rpoC:gfp SpR* | This study |
| *ΔspoIIIAH* | *Bacillus subtilis subsp. subtilis trpC2 ΔspoIIIAH::erm* | Koo *et al.* (40) |
| *ΔspoIIQ* | *Bacillus subtilis subsp. subtilis trpC2 ΔspoIIQ::erm* | Koo *et al.* (40) |
| *ΔspoIID* | *Bacillus subtilis subsp. subtilis trpC2 ΔspoIID::erm* | Koo *et al.* (40) |
| *ΔspoIIIAH* RpsB-GFP | *Bacillus subtilis subsp. subtilis trpC2 ΔspoIIIAH::erm rpsB:gfp SpR* | This study |
| *ΔspoIIQ* RpsB-GFP | *Bacillus subtilis subsp. subtilis trpC2 ΔspoIIQ::erm rpsB:gfp SpR* | This study |
| *ΔspoIIQ* RpoC-GFP | *Bacillus subtilis subsp. subtilis trpC2 ΔspoIIQ::erm rpoC:gfp SpR* | This study |
| *ΔspoIID* RpsB-GFP | *Bacillus subtilis subsp. subtilis trpC2 ΔspoIID::erm rpsB:gfp SpR* | This study |
| *ΔspoIID* RpoC-GFP | *Bacillus subtilis subsp. subtilis trpC2 ΔspoIID::erm rpoC:gfp SpR* | This study |

**B. Primers**

| Primer | Sequence |
|---|---|
| ***rpsB*** | |
| UPFOR | GATACCTACGCCTCGTTTAGAATTCGCGGCGCAATC |
| UPREV | CATTGATCCGCTGCCTGATCCGGACGCAGTTGTTGTTTCTGTTTC |
| MID1FOR | GAAACAGAAACAACAACTGCGTCCGGATCAGGCAGCGGATCAATG |
| MID1REV | GTATTTTTCCGTTAATCAAATTGCTCATTCACTTATAGAGTTCATCCATACC |
| MID2FOR | GGTATGGATGAACTCTATAAGTGAATGAGCAATTTGATTAACGGAAAAATAC |
| MID2REV | GTCCCTCTTATCACCTTTTGAATAGGTAATTGAGAGAAGTTTCTATAGAATTTTTC |
| DOWNFOR | GAAAAATTCTATAGAAACTTCTCTCAATTACCTATTCAAAAGGTGATAAGAGGGAC |
| DOWNREV | GTGTCTGCGCTCCGTAATATTCAACCGTTACTTTATCTAATAATG |
| ***rpoC*** | |
| UPFOR | CTGATGTTAAAGCTGTTATCGCACAGCAGC |
| UPREV | CATTGATCCGCTGCCTGATCCGGATTCAACCGGGACCATATCGTCAG |
| MID1FOR | CTGACGATATGGTCCCGGTTGAATCCGGATCAGGCAGCGGATCAATG |
| MID1REV | GTATTTTTCCGTTAATCAAATTGCTCATTCACTTATAGAGTTCATCCATACC |
| MID2FOR | GGTATGGATGAACTCTATAAGTGAATGAGCAATTTGATTAACGGAAAAATAC |
| MID2REV | CAGTCTTTCAGCAGAGTTAAATCAGTAATTGAGAGAAGTTTCTATAGAATTTTTC |
| DOWNFOR | GAAAAATTCTATAGAAACTTCTCTCAATTACTGATTTAACTCTGCTGAAAGACTG |
| DOWNREV | ATGAGCGTTTGCTTGAAGACGGTCTCTTAAAG |

**C. Plasmids**

| Plasmid | Description |
|---|---|
| pSHP2 (48) | Used as a template for GFP and spectinomycin resistance cassette |

### References and Notes


1. D. Z. Rudner, R. Losick, Protein Subcellular Localization in Bacteria. *Cold Spring Harb. Perspect. Biol.* **2**, a000307 (2010).

**Movie S1. Timelapse of WT RpsB-GFP cells during sporulation.**

**Movie S2. Timelapse of WT RpoC-GFP cells during sporulation.**

**Data S1. Mass spectrometry results**

The data, deposited in ProteomeXchange (PXD047497), were processed with MaxQuant, and log2 fold changes for 10 experimental groups vs Ctrl were calculated, revealing insights into the ribosomal proteome dynamics during sporulation.
